# Computational analysis of ligand dose range thermal proteome profiles

**DOI:** 10.1101/2020.05.08.083709

**Authors:** Nils Kurzawa, Isabelle Becher, Sindhuja Sridharan, Holger Franken, André Mateus, Simon Anders, Marcus Bantscheff, Wolfgang Huber, Mikhail M. Savitski

## Abstract

Detecting ligand-protein interactions in living cells is a fundamental challenge in molecular biology and drug research. Proteome-wide profiling of thermal stability as a function of ligand concentration promises to tackle this challenge. We present a statistical analysis method with reliable control of the false discovery rate and apply it to several datasets probing epigenetic drugs. We detect off-target drug engagement in unrelated protein families.

## Main

Studying ligand-protein interactions is essential for understanding drug mechanisms of action and adverse effects (Becher et al., 2016; Savitski et al., 2014), and more generally for gaining insights into molecular biology by enabling the monitoring of metabolite- and protein-protein interactions (Dai et al., 2018; Piazza et al., 2018; Sridharan et al., 2019; Tan et al., 2018, Becher et al. (2018)). Thermal proteome profiling (TPP) (Franken et al., 2015; Mateus et al., 2020; Savitski et al., 2014) combines quantitative, multiplexed mass spectrometry (MS) (Bantscheff et al., 2012) with the cellular thermal shift assay (Martinez Molina et al., 2013) and enables proteome-wide measurements of thermal stability, by quantifying non-denatured fractions of cellular proteins across temperatures. TPP has been used to study binding of ligands and their downstream effects in cultured human (Becher et al., 2016; Savitski et al., 2014) and bacterial cells (Mateus et al., 2018), and has recently been adapted to animal tissues and human blood (Perrin et al., 2020).

Besides temperature, non-denatured fractions of cellular proteins can be measured as a function of other variables, such as ligand concentration. In the 2D-TPP experimental design, both temperature and ligand concentration are systematically varied (Becher et al., 2016). In comparison to the original temperature range TPP (TPP-TR), this method overcomes the problem that different proteins may be susceptible to thermal stability modulation at different compound concentrations or temperatures. Thus, 2D-TPP can greatly increase sensitivity and coverage of the amenable proteome. However, while statistical analysis for the TPP-TR assay is well established (Childs et al., 2019; Franken et al., 2015), similar approaches for 2D-TPP have been hampered by its more complicated experimental design. 2D-TPP employs a multiplexed MS analysis of samples in the presence of *n* ligand concentrations (including a vehicle control) at *m* temperatures (Supplementary Figure 1). Thus, for each protein *i*, a *m* x *n* data matrix *Y*_*i*_ of reporter ion intensities is observed. However, these matrices contain non-randomly missing values, usually at higher temperatures, due to differential thermal stability across the proteome, i.e., some proteins may fully denature at one of the temperatures used in the experiment and thus will not be quantified at this or higher temperatures.

In the approach of Becher et al. (2016), non-linear dose-response curves were fitted to each protein for each individual temperature. Subsequently, hits were defined by applying bespoke rules, including a requirement for two dose-response curves at consecutive temperatures to both have *R*^2^ > 0.8 and a fold change of at least 1.5 at the highest treatment concentration. However, this approach, with its reliance on data-independent thresholds, has uncontrolled specificity (i.e., there is no possibility for controlling the false discovery rate (FDR)), and as a consequence, may have suboptimal sensitivity if, e.g., the thresholds are too stringent.

Here, we present a statistical method for FDR-controlled analysis of 2D-TPP data. Our approach fits two nested models to protein reporter ion intensities. The null model allows the soluble protein fraction to depend on temperature, but not concentration, as expected for proteins with no treatment induced change in thermal stability. The alternative model fits the soluble protein fraction as a sigmoid dose-response function of concentration, separately for each temperature. To increase robustness, certain parameters of the sigmoids are shared or constrained across temperatures: the slope and the direction of the response (destabilization or stabilization) are set to be the same across temperatures, and the inflection point (EC50) is required to increase or decrease linearly with temperature. The residual sum of squares (RSS) of the two models are compared to obtain, for each protein, an *F*-statistic. To calibrate this statistic in terms of FDR, we adapted the bootstrap approach of Storey et al. (2005). Briefly, residuals from the alternative model are resampled and added back to the null model estimate to simulate the case where there is no concentration effect of the ligand on the thermal profile of a protein. The resampling scheme takes into account the noise dependence of measurements within individual MS runs. Overall, our approach allows the detection of ligand-protein interactions from thermal profiles (DLPTP; Fig. 1a; Supplementary Figure 2).

**Figure 1:**
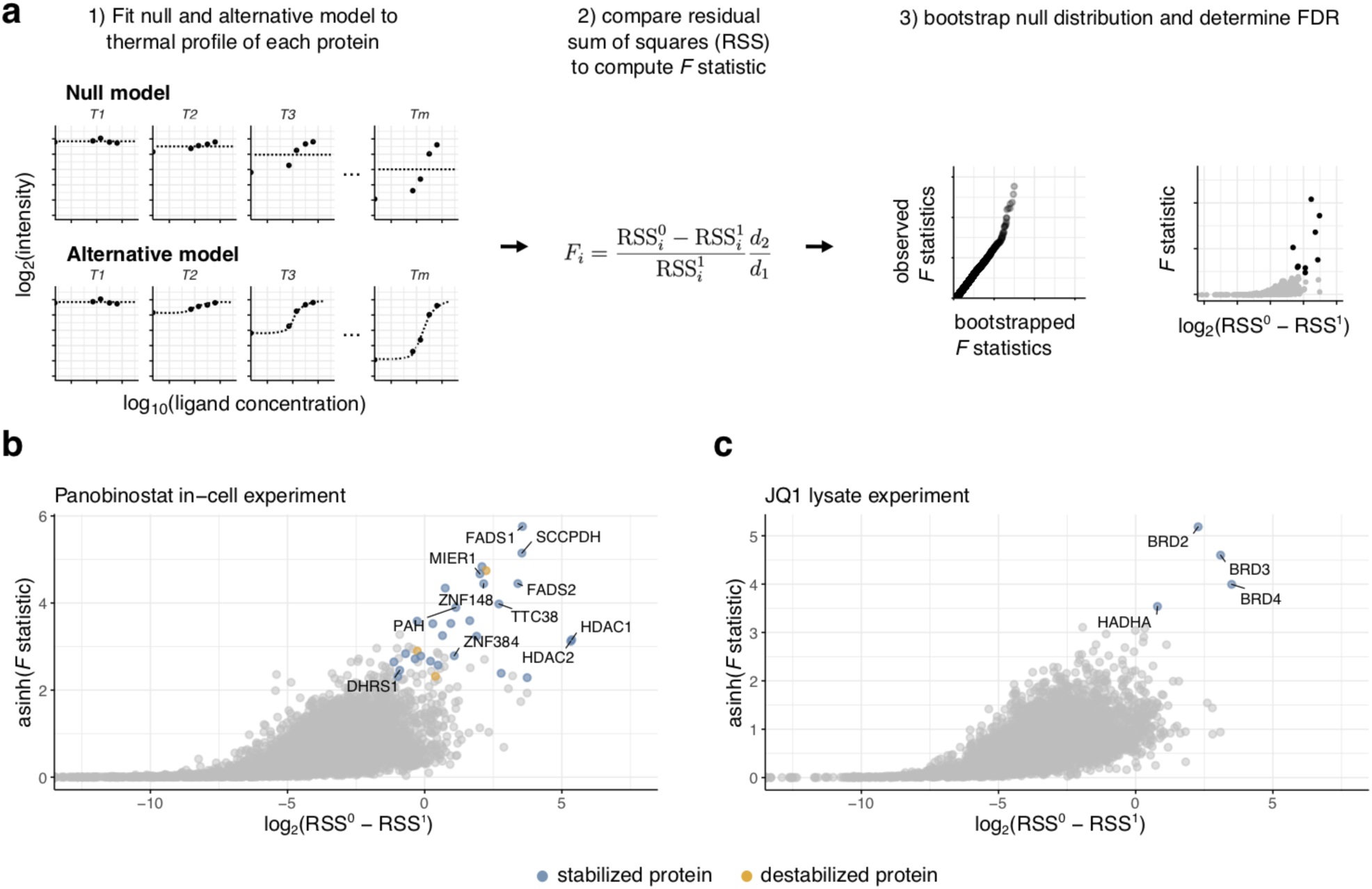
DLPTP recovers drug-protein interactions. **a** Illustration of fitted curves under the null and alternative model and how obtained residuals are used to find proteins significantly altered in their thermal stability (example fits for the null and alternative model are shown in Supplementary Figure 2). **b** Volcano plot for the 2D-TPP dataset acquired upon treatment of HepG2 cells with the HDAC-inhibitor panobinostat. **c** Analogous to **b**, for the JQ1 2D-TPP dataset acquired in THP1 lysate. **b, c** Blue and orange points represent proteins that were detected as stabilized or destabilized respectively by the drug treatment, at 10% FDR.

We applied our approach by re-analysing previously published 2D-TPP datasets. For the pan-HDAC inhibitor panobinostat (Laubach et al., 2015) profiled in intact HepG2 cells, we recovered all previously reported on- and off-targets of the drug (Becher et al., 2016): HDAC1, HDAC2, TTC38, PAH, FADS1 and FADS2 (Fig 1b; Supplementary Figure 3a and c; Supplementary Data 1), except HDAC6, which showed a noisy, non-sigmoidal profile in this dataset (Supplementary Figure 3e; Supplementary Data 1). In addition, we found more recently discovered interactors of panobinostat: DHRS1 and two zinc-finger transcription factors ZNF148 and ZNF3841 (Perrin et al., 2020) that had not been found by previous analyses of this dataset.

Next, we re-analyzed a dataset probing the BET bromodomain inhibitor JQ1 in THP1 cell lysate (Savitski et al., 2018). We recovered previously reported targets BRD2, 3, 4 and HADHA, an enzyme with acetyltransferase activity, (Fig 1c; Supplementary Figure 3b and d; Supplementary Data 1). These analyses showcase that DLPTP can be applied to 2D-TPP experiments acquired in intact cells as well as in lysates.

Next, we performed a 2D-TPP experiment in HL60 cells with the epigenetic inhibitor PCI-34051 (Supplementary Figure 4a), a reported selective inhibitor of HDAC8 which was suggested as a potential treatment for multiple types of T-cell leukemia (Balasubramanian et al., 2008). Apart from an enrichment of proteins annotated for ‘oxidation-reduction process’ and ‘carboxylic acid metabolic process’ showing significant thermal stability effects (hypergeometric test, adjusted *p* = 6.32 · 10^−6^ and 6.70 · 10^−5^, respectively), likely reflecting the cellular response to the drug treatment, we found HDAC8 (pEC_50,PCI-34051_ = 6.40) as one of the top stabilized hits (Fig. 2a; Supplementary Figure 5a). Interestingly, among the significantly affected proteins by PCI-34051 we also found Leucine aminopeptidase 3 (LAP3). The expression of LAP3 correlates with tumor cell proliferation (Tian et al., 2014), and its inhibition suppresses invasion of ovarian cancer (Wang et al., 2015). To follow-up our identification of LAP3, we turned to BRD-3811, an analog of PCI-34051 (Supplementary Figure 4b), in which the Zn^2+^ chelating hydroxamic acid (HA) group is sterically hindered from binding HDAC8 by an additional methyl group. A 2D-TPP experiment in HL60 cells with BRD-3811 showed no significant stabilization of HDAC8, as expected. However, we again found LAP3 as a significant target (Fig. 2b; Supplementary Figure 5b). To investigate whether LAP3 function was inhibited by binding of either version of the compound, we performed an *in vitro* fluorometric leucine aminopeptidase assay using a recombinant LAP3 enzyme. Indeed, we found that both compounds inhibited recombinant LAP3 peptidase activity *in vitro* (Fig. 2c). However, BRD-3811 showed a reduced effect, in line with a 10-fold lower pEC_50_ measured in the 2D-TPP experiment (pEC_50,PCI-34051_ = 5.94, pEC_50,BRD-3811_ = 4.98), which suggests that the binding of both molecules to LAP3 might be mediated via the HA group, but dampened by the additional methyl group in the case of BRD-3811. In conclusion, 2D-TPP of the analog compounds PCI-3405 and BRD-3811 and DLPTP analysis revealed their intracellular target space (Supplementary Data 2) and showed that both bind and inhibit LAP3, a potentially interesting cancer target.

**Figure 2:**
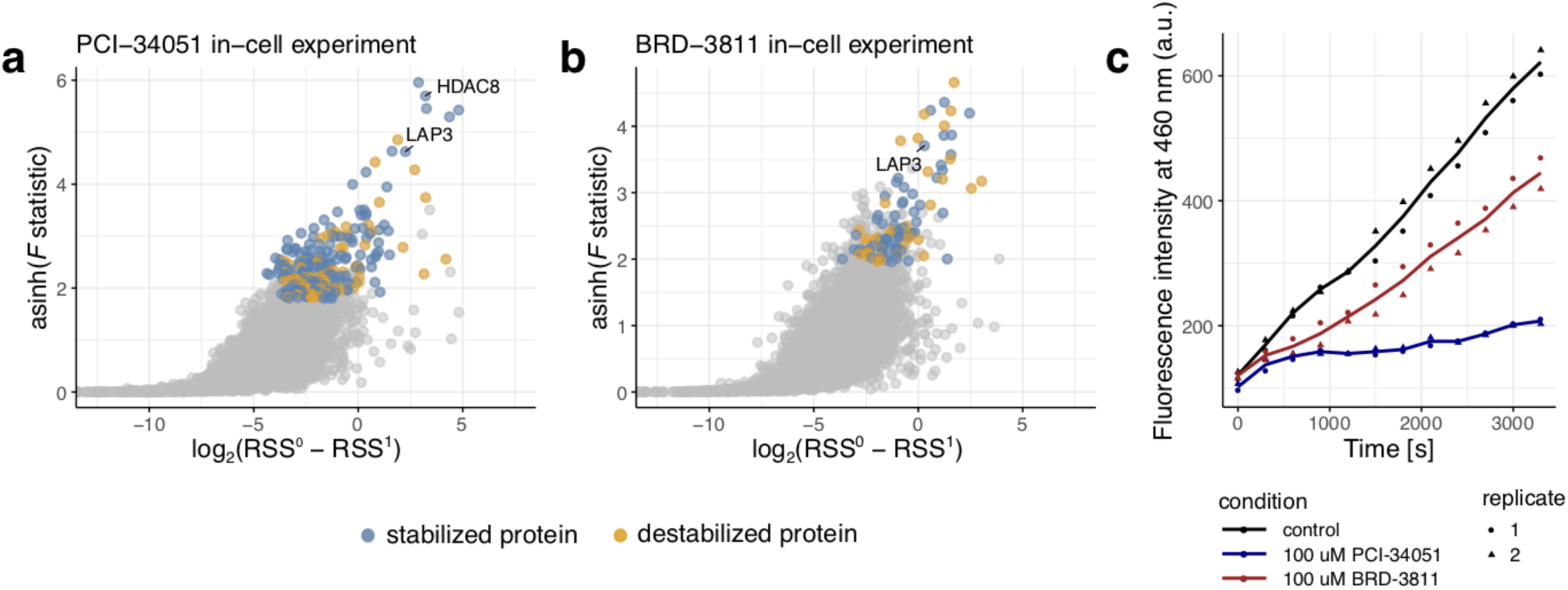
DLPTP reveals LAP3 as a target of PCI-34051 with and without the hydroxamic acid group. **a** Volcano plot of the 2D-TPP experiment with PCI-34051 in HL-60 cells. **b** Volcano plot of the 2D-TPP experiment with BRD-3811 in HL-60 cells. **a, b** Blue and orange points correspond to significant stabilization and destabilization hits at 10% FDR, **c** Fluorescence intensity measured over time in a fluorometric leucine aminopeptidase assay with recombinant LAP3 in the presence of PCI-34051, BRD-3811 or vehicle control.

In summary, we describe DLPTP, a statistical method for analysis of 2D-TPP datasets with FDR control via bootstrap resampling, which is available as an open-source R package from Bioconductor (https://bioconductor.org/packages/TPP2D). We exemplify the use of our approach by applying it to previously published and novel 2D-TPP datasets. First, our reanalysis of panobinostat and JQ1 datasets identified the cognate targets, as well as previously described off-targets (Becher et al., 2016; Perrin et al., 2020; Savitski et al., 2018). Second, we discovered the engagement of LAP3 by the HDAC8 inhibitor PCI-34051 and its analog BRD-3811 and found that both interactions lead to inhibition of LAP3 peptidase activity in an *in vitro* assay. This finding opens possibilities towards the development of specific LAP3 inhibitors based on the structure of BRD-3811.

We focus here on studies of high-affinity ligands such as drugs and tool compounds. However, DLPTP is also applicable for the detection of more transient ligand-protein interactions, as shown, e.g., in an analysis of a 2D-TPP dataset of gel-filtered Jurkat cell lysate treated with GTP. We found a strong enrichment of GTP and ATP binding annotated proteins among the detected hits (Supplementary Note Figure 2; Supplementary Data 3; details on the analysis in the Supplementary Note), in line with recent reports (Piazza et al., 2018; Sridharan et al., 2019). We believe that our approach will improve sensitivity for detecting ligand-protein interactions from thermal proteome profiling experiments with drugs and other ligands and will help prevent expense of effort on false positives.

## Methods

### 2D-TPP Experiments

2D-TPP experiments for profiling PCI-34051 and BRD-3811 were performed as described (Becher et al., 2016). Briefly, HL-60 (DSMZ) cells were grown in IMDM medium supplied with 10% FBS. Cells were treated with a concentration range (0, 0.04, 0.29, 2, 10 *µ*M) of PCI-34051 (Selleckchem) or BRD-3811 (prepared as described previously (Olson et al., 2014)) for 90 min at 37 °C, 5% CO_2_. The samples from each treatment-concentration were split into 12 portions which were then heated over a temperature range (42 - 63.9 °C) for 3 min, further sample processing was performed as described (Becher et al., 2016).

The 2D-TPP experiment to assess GTP binding proteins was performed using gel filtered lysate as described (Becher et al., 2016). In short, JurkatE6.1 cells were cultured in RPMI (GIBCO) medium supplemented with 10% heat inactivated FBS. The cells were harvested and washed with phosphate buffered saline (PBS - 2.67 mM KCl, 1.5 mM KH_2_PO_4_, 137 mM NaCl, and 8.1 mM NaH_2_PO_4_, pH 7.4). The cell pellet was resuspended in lysis buffer (PBS containing protease inhibitors and 1.5 mM MgCl_2_) equal to 10 times the volume of the cell pellet. The cell suspension was lysed by mechanical disruption using a Dounce homogenizer (20 strokes) and treated with benzonase (25 U/ml) for 60 min at 4°C on a shaking platform. The lysate was ultracentrifuged at 100,000 g, 4°C for 30 min. The supernatant was collected and deslated using PD-10 column (GE-healthcare). The protein concentration of the eluted lysate was measured using Bradford assay. The protein concentration of the lysate was maintained at 2 mg/ml for the assay. The lysate was treated using a concentration-range of GTP (0, 0.001, 0.01, 0.1, 0.5 mM) for 10 min at room temperature. The samples from each GTP-concentration were split into 12 portions which were then heated over a temperature range (42 - 63.9 °C) for 3 min. Post-heat treatment, the protein aggregates were removed using ultracentrifugation at 100,000 g, 4°C for 20 min. Subsequently, the supernatants were processed as previously described (Becher et al., 2016).

### Protein identification and quantification

Raw mass spectrometry data were processed with Isobarquant (Franken et al., 2015) and searched with Mascot 2.4 (Matrix Science) against the human genome (FASTA file downloaded from Uniprot, ProteomeID: UP000005640) extended by known contaminants and reversed protein sequences (search parameters: trypsin; missed cleavages 3; peptide tolerance 10 ppm; MS/MS tolerance 0.02 Da; fixed modifications were carbamidomethyl on cysteines and TMT10-plex on lysine; variable modifications included acetylation on protein N-terminus, oxidation of methionine, and TMT10-plex on peptide N-termini). Protein FDR was calculated using the picked approach (Savitski et al., 2015).

Reporter ion spectra of unique peptides were summarized to the protein level to obtain the quantification *s*_*i,u*_ for protein *i* measured in condition *u* = (*j, k*), i.e., at temperature *j* and concentration *k*. Isobarquant additionally computes robust fold changes between the different treatment conditions *r*_*i,u*_ for each protein *i* in condition *u* relative to control condition *u*′ using a bootstrap approach. We used these to obtain per-condition log_2_ signal intensities computed as *y*_*i,u*_ = log_2_((*r*_*i,u*_/ Σ_*l*_ *r*_*i,l*_) Σ_*l*_ *s*_*i,l*_). This step is not mandatory, and there is no reason our method could not also be used with input from other quantification softwares.

From the obtained abundance tables, proteins in condition *u* were removed that did not fulfil the criterion of having been quantified by more than one unique peptide and a total number of measurements *p*_*i*_ per protein *i* of at least 20 was required.

The MS experiment comprising the temperatures 54 and 56.1°C was removed from the PCI-34051 dataset for analysis as we noticed that it contained unexpectedly high noise levels. In particular the relative reporter ion intensities at 54°C showed about ten times higher variance than all other temperatures, likely due to a drop in instrument performance during the time this sample was measured.

Moreover, we noted that measured profiles of some proteins appeared to have been affected by carry over from previous experiments for the PCI-34051 and BRD-3811 datasets. These profiles exhibited a characteristic pattern as depicted in Supplementary Figure 5c in which apparent stabilization of these proteins was observed only in half of the TMT channels corresponding to every other temperature. These proteins were filtered out by manual inspection.

### Data pre-processing of public datasets

The panobinostat and JQ1 datasets were downloaded from the publisher websites as spreadsheets that were provided as supplementary data together with the publications (Becher et al., 2016; Savitski et al., 2018). Log_2_ signal intensities were computed as described above and retrieved tables were filtered such that each protein contained at least one measurement quantified by more than one unique peptide and *p* ≥ 20 observations.

### Model description

Two nested models were fitted to the thermal profile of each protein *i*. We modeled observed intensity values of protein *i* at a given temperature *j* and ligand concentration *k*. In case of the null model we assumed:

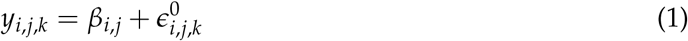

Here, the base intensity level at a given temperature *j* is captured by *β*_*i,j*_. For the alternative model we assumed:

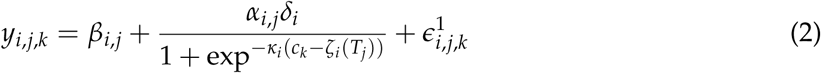

Here, the base intensity level at a given temperature *j* is captured by *β*_*i,j*_j, *δ*_*i*_ describes the maximal absolute stabilization across all temperatures, *α*_*i,j*_ ∈ [0, 1] indicates how much of the maximal stabilization occurs at temperature *j* and *k*_*i*_ is a common slope factor fitted across all temperatures. Finally, *ζ*_*i*_(*T*_*j*_) is the concentration of the half-maximal stabilization, with 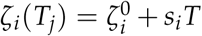, where *s*_*i*_ is a slope representing a linear temperature-dependent decay or increase of the inflection point and 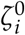 is the intercept of the linear model. Based on the residual sum of squares (RSS): 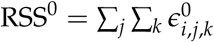 and 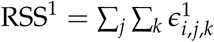 obtained from the two fitted models, we computed an *F* statistic:

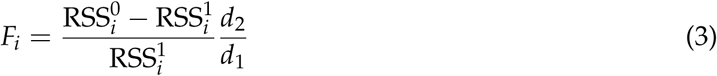

with the degrees of freedom *d*_1_ = *v*_1_ −*v*_0_ and *d*_2_ = *p*_*i*_ − *v*_1_ where *p*_*i*_ is the number of observations for protein *i* that were fitted, and *v*_0_ and *v*_1_ are the number of parameters of the null and alternative model respectively.

### FDR estimation

In order to estimate the false discovery rate (FDR) associated with a given threshold *θ* for the *F* statistic obtained for a protein *i* with *m*_*i*_*n*_*i*_ number of observations, we adapted the bootstrap approach of Storey et al. (2005) as follows. To generate a null distribution, the following was repeated *B* times: i) Resample with replacement the residuals 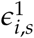 obtained from the alternative model fit for protein *i* in MS experiment *s* to obtain 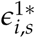 and add them back to the corresponding fitted estimates of the null model to obtain 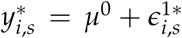. ii) Fit null and alternative models to 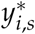 and compute the *F*-statistic *F*^0*b*^. An FDR was then computed by partitioning the set of proteins {1, …, *P*} into groups of proteins with similar number *D*(*p*) of measurements, e.g. *γ* (*p*)= ⌊*D*(*p*)+ 1⌋ and then

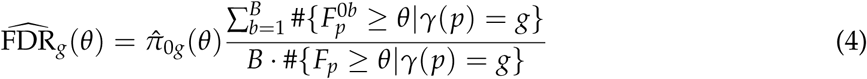

The proportion of true null events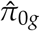 in the dataset of proteins in group *g* was estimated by:

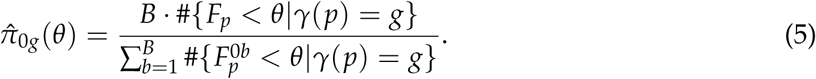

### Fluorometric aminopeptidase assay

LAP3 activity was determined using the Leucine Aminopeptidase Activity Assay Kit (abcam, ab124627) and recombinant LAP3 (origene, NM_015907). Recombinant LAP3 enzyme was dissolved in the kit assay buffer and incubated for 10 minutes at room temperature with vehicle (DMSO) or 100 *µ*M of either PCI-34051 or BRD-3811. All other assay steps were performed as described in the kit. Fluorescent signal (Ex/Em = 368/460 nm) was detected over 55 minutes.

## Code availability

The software is free and open source as an R package from Bioconductor: https://bioconductor.org/packages/TPP2D

## Data availability

All acquired mass spectrometry datasets (2D-TPP experiment of GTP treated gel-filtered Jurkat lysate, PCI-34051 and BRD-3811 treated HL-60 cells) were deposited on PRIDE (accession number: PXD016640).

## Acknowledgements

The authors thank Srishti Dar, Henrik Hammarén, Britta Velten, Carola Doce, Dorothee Childs, Toby Mathieson and Stephan Gade for insightful discussions and critical feedback. This work was supported by the European Molecular Biology Laboratory (EMBL). NK was supported by a fellowship of the EMBL International PhD programme. SA is funded by the Deutsche Forschungsgemeinschaft, SFB 1036. WH acknowledges funding from the European Commission’s H2020 Programme, Collaborative research project SOUND (Grant Agreement no 633974).

## Author contributions

NK, IB, MB, WH and MMS conceived the project, designed experiments and outlined desired method and software features. NK implemented and applied the software and performed data analysis. IB and SS performed experiments. HF and AM benchmarked and evaluated the method and gave input. SA gave crucial input. MB, WH and MMS jointly supervised the work. NK wrote the manuscript with feedback from all authors.

## Supplementary Figures

**Supplementary Figure 1:**
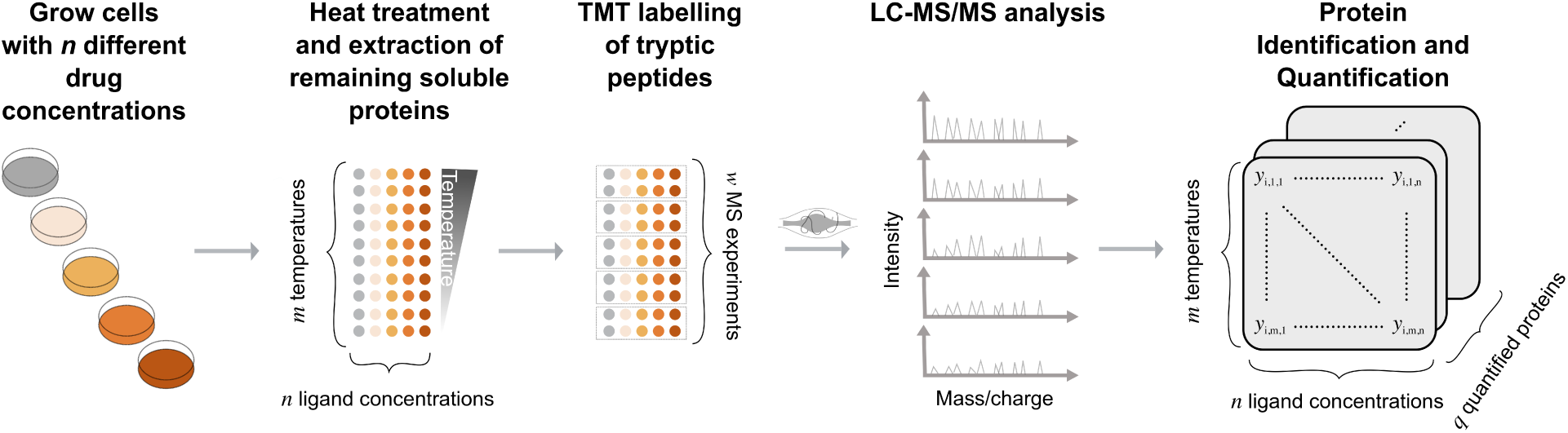
Illustration of the 2D-TPP experimental setup. Cells are grown in the presence of *n* different concentrations of a ligand of interest. Each sample is divided into *m* aliquots, each of which is subjected to one of m temperatures, and the remaining soluble proteins are extracted. Proteins are digested with trypsin and labelled with TMT, such that one set of TMT labels is used for all concentrations and two adjacent temperatures. *w* = *m*/2 MS runs are performed, and peptides are identified by database search and quantified signal is aggregated on the protein level.

**Supplementary Figure 2:**
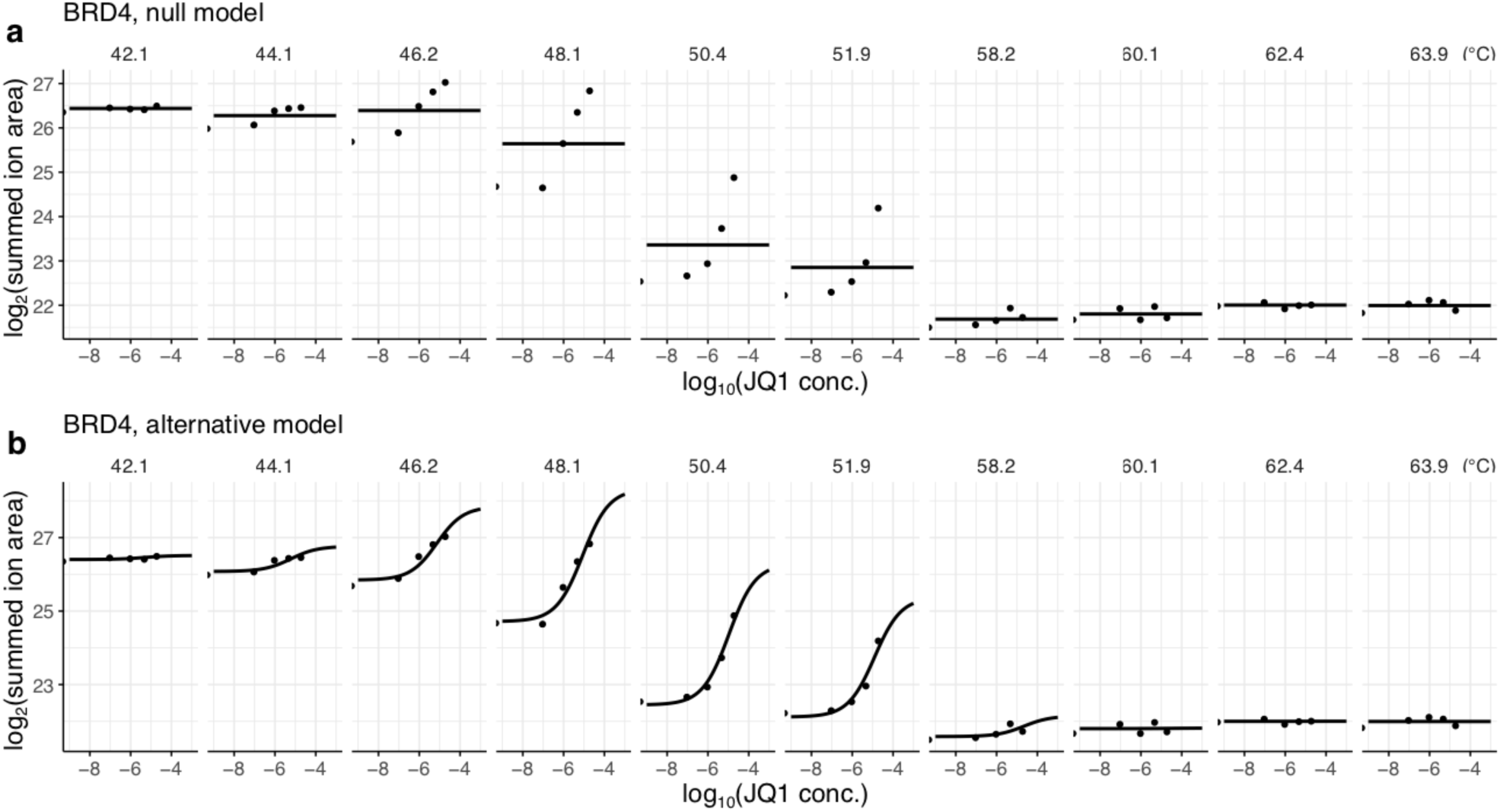
Example of null and alternative model fits to the thermal profile of BRD4 measured in THP1 lysate in the presence of different concentrations of its inhibitor JQ1.

**Supplementary Figure 3:**
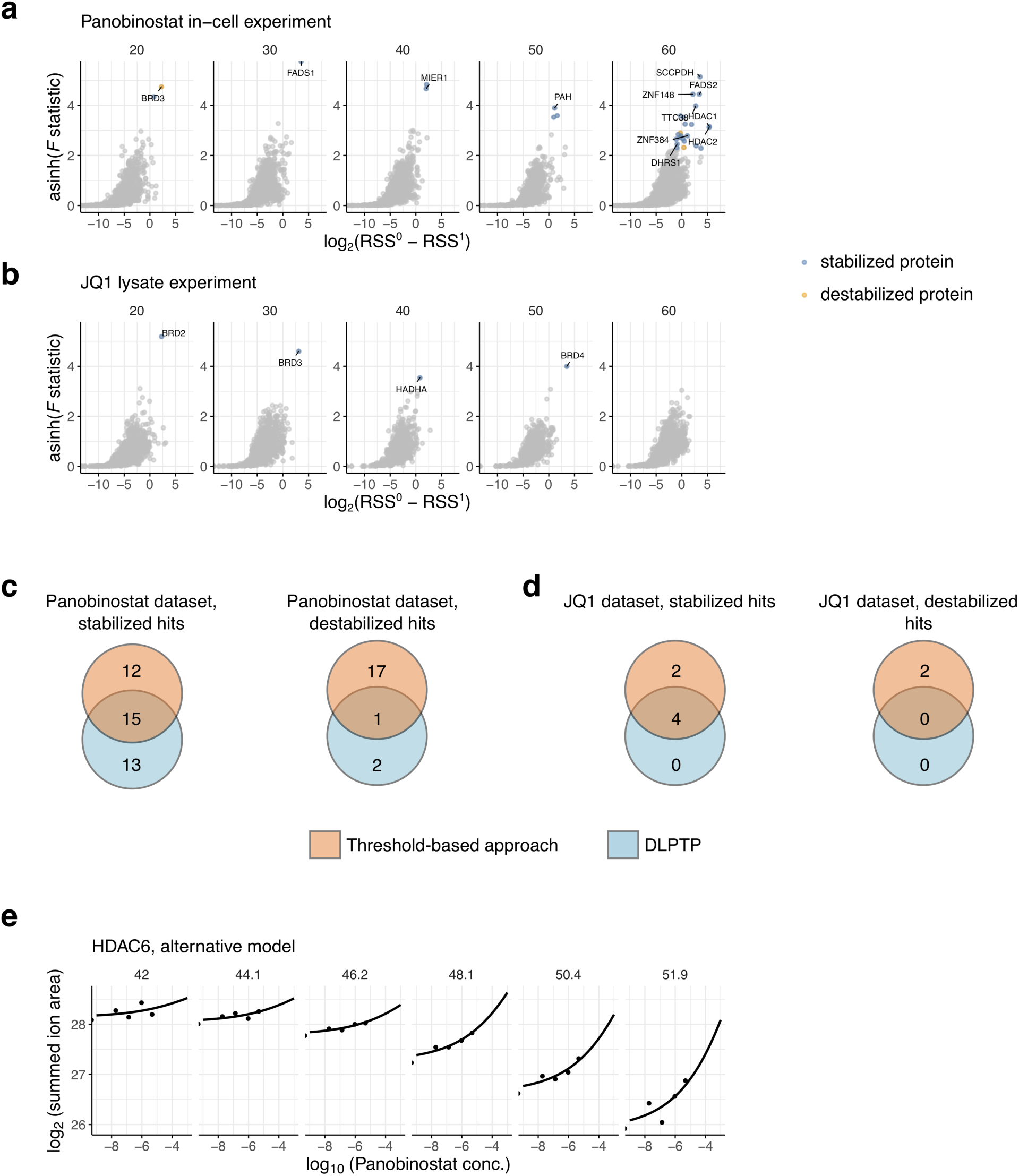
Comparison of 2D-TPP analysis results obtained by DLPTP and the threshold-based approach. **a** Volcano plot for the 2D-TPP experiment of panobinostat acquired in HepG2 cells of proteins grouped by similar number of observations, e.g., 30 corresponds to 24 < *p* < 35. **b** Same as **a** for the 2D-TPP experiment of JQ1 in THP1 lysate. **c** Venn diagrams comparing hits found in the panobinostat dataset between DLPTP and the threshold-based approach. **d** same as **c** for the JQ1 dataset. **e** Alternative model fit for HDAC6 in the Panobinostat dataset showing that the protein has a noisy, non-sigmoidal stabilization profile.

**Supplementary Figure 4:**
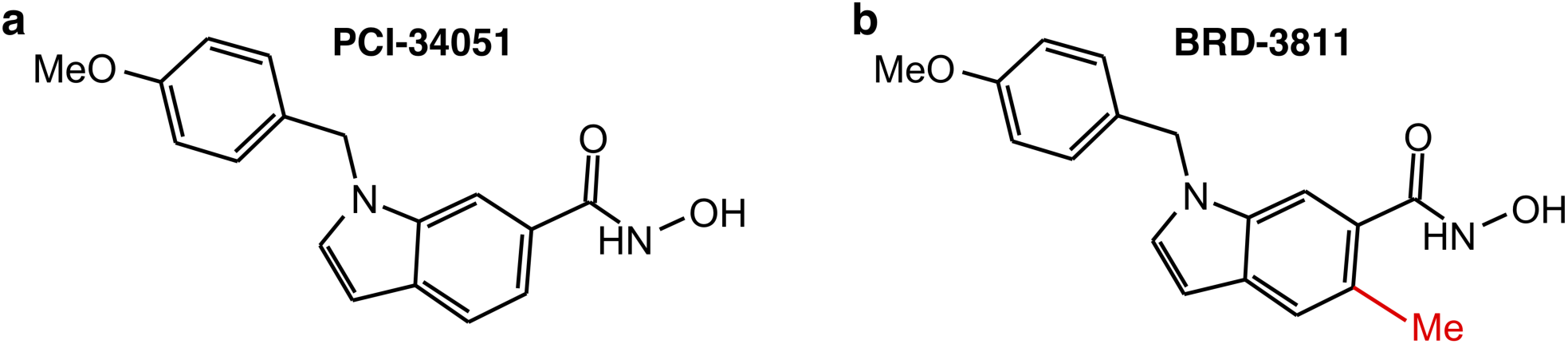
Chemical structures of PCI-34051 and BRD-3811.

**Supplementary Figure 5:**
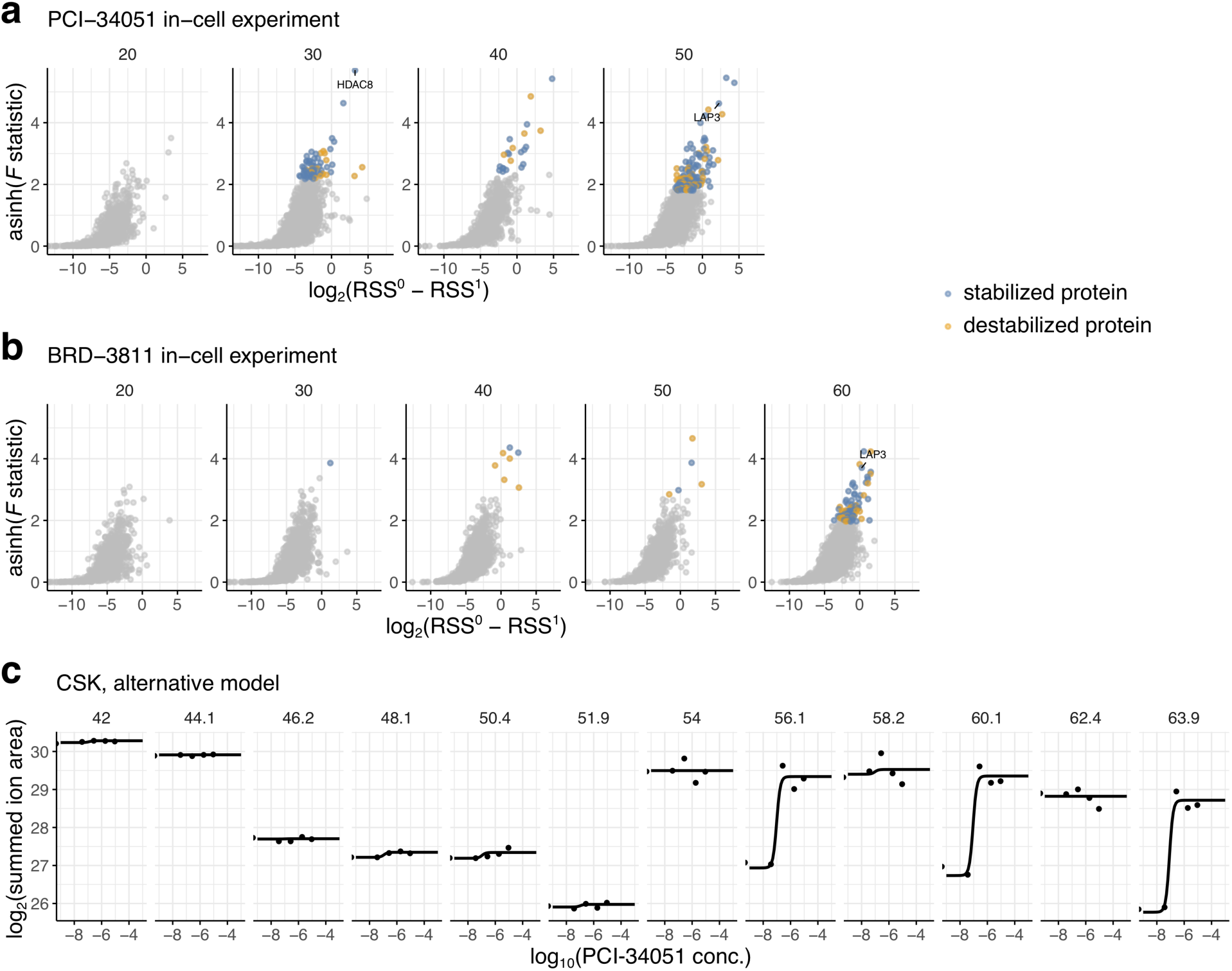
Volcano plots grouped by proteins with similar number of observations and example of thermal profile of CSK affected by carry-over. **a** Volcano plot for the 2D-TPP experiment of PCI-34051 acquired in HL-60 cells of proteins grouped by similar number of observations, e.g., 30 corresponds to 24 < p < 35. **b** Same as **a** for the 2D-TPP experiment of BRD-3811 in HL-60 cells. **c** 2D thermal profile of Casein kinase (CSK) obtained in the dataset of PCI-34051 profiled in HL-60 cells.

## Supplementary Note

### Evaluation of the implementation’s FDR-control

To evaluate whether our method implementation controlled FDR as expected and how its sensitivity compared to the threshold-based approach, we created a synthetic dataset for benchmarking. It was composed of 5000 simulated protein thermal profiles expected under the null hypothesis with gaussian noise based on standard deviations observed for real datasets and 80 spiked-in known true positive protein profiles obtained from various different datasets. We then ran our method on this dataset using *B* = {5, 20, 100} rounds of bootstrapping to generate null distributions. We found that our method faithfully controlled FDR at 1, 5 and 10% for *B* = {20, 100} and that its sensitivity was higher than the threshold-based approach (Supplementary Note Figure 1).

**Supplementary Note Figure 1:**
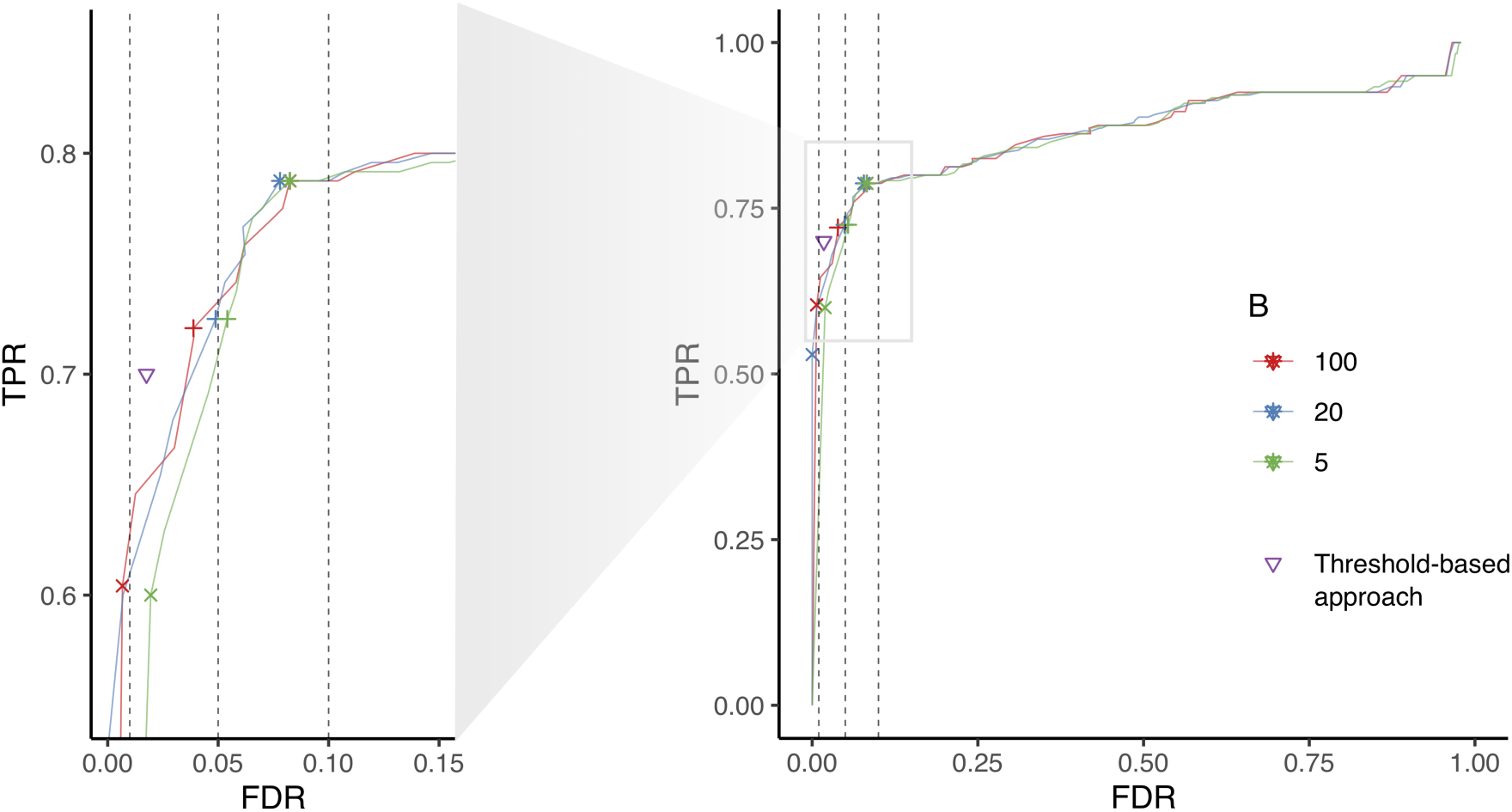
Benchmarking of DLPTP on synthetic dataset confirms FDR-control. True positive rate (TPR, i.e. sensitivity) versus FDR curves of DLPTP’s performance when bootstrapping the null distribution with *B* = {5, 20, 100}. Crosses (x) represent results for FDR set to 1%, pluses for 5%, asterisks (*) for 10% and the triangle represents performance of the threshold-based approach suggested by Becher et al. (2016). Data shown for DLPTP’s performance for different *B* is an average of n = 3.

### Analysis of the GTP 2D-TPP experiment in gel filtered Jurkat lysate

To showcase the application of DLPTP to a 2D-TPP dataset profiling metabolite-protein interactions, we performed a 2D-TPP experiment in gel filtered lysate of Jurkat cells treated with a concentration range from 0 to 0.5 mM of NaGTP. The gel filtration leads to a depletion of endogenous metabolites and thus makes metabolite-interacting proteins, which often otherwise remain strongly bound to metabolites in lysates, particularly susceptible to bind to externally supplied metabolites.

Among the significantly stabilized proteins we found when applying DLPTP to this dataset, proteins annotated with the Gene Ontology terms ‘GTP binding’ (hypergeometric test *p* < 2.2 · 10^−16^, odds ratio: 19.85) and ‘ATP binding’ (hypergeometric test *p* < 6.81 · 10^−9^, odds ratio: 3.26) were significantly enriched (Supplementary Note Figure 2a; Supplementary Data 3). In comparison to the threshold-based approach DLPTP recovered more annotated GTP and ATP binders at 10% FDR, whereas at 1% FDR it found fewer false positives, i.e. proteins not annotated for GTP or ATP binding (Supplementary Note Figure 2b).

**Supplementary Note Figure 2:**
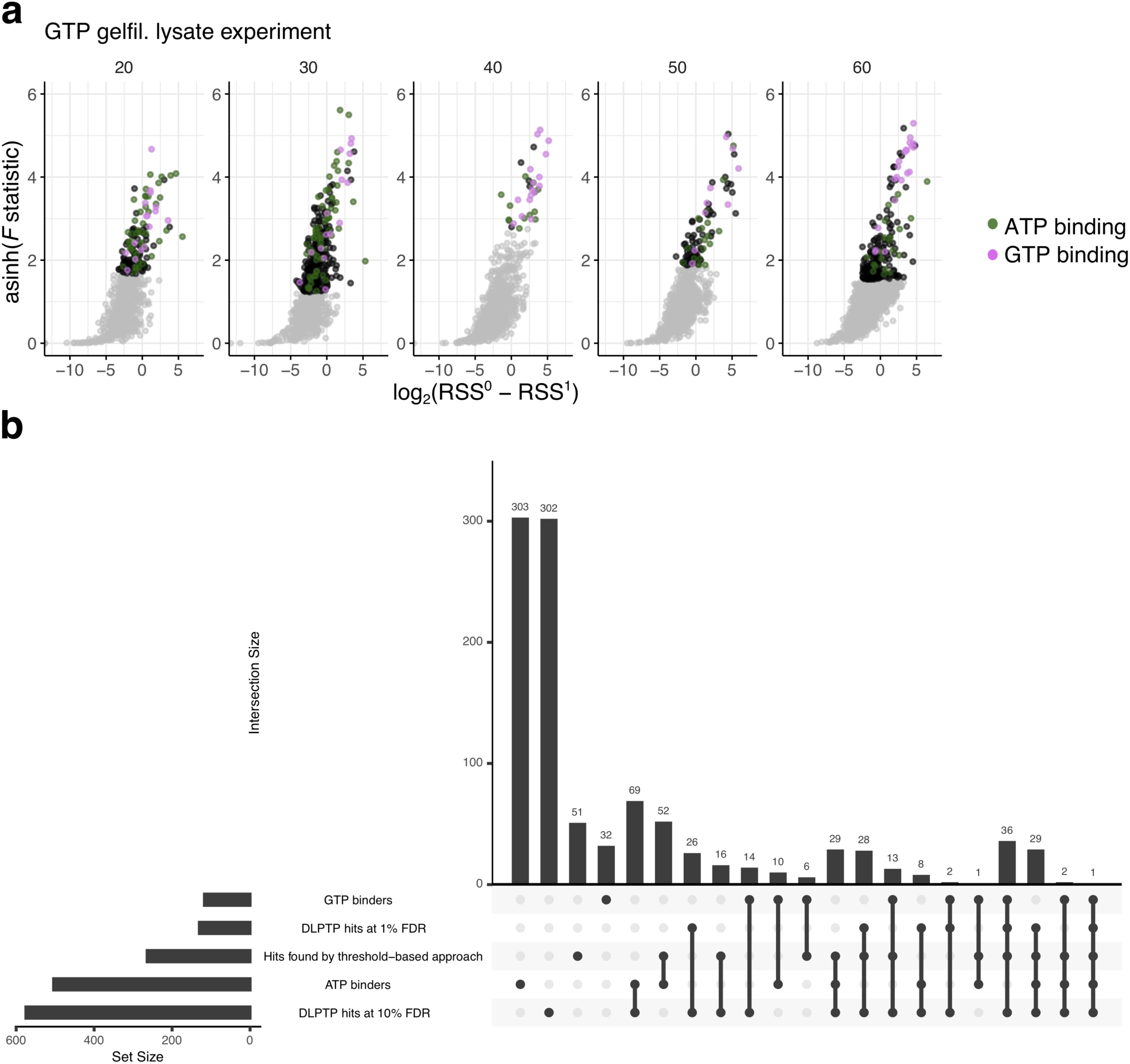
DLPTP recovers annotated GTP and ATP interactors from a 2D-TPP experiment profiling GTP-protein interactions. **a** Volcano plot for the analysis of the GTP gel-filtered lysate dataset at 10% FDR. **b** Upset plot of set intersections of annotated GTP and ATP-binding proteins, hit found with the threshold-based method by Becher et al.

